# A midbrain dynorphin circuit promotes threat generalization

**DOI:** 10.1101/2020.04.21.053892

**Authors:** Lizz Fellinger, Yong S. Jo, Avery C. Hunker, Marta E. Soden, Larry S. Zweifel

## Abstract

Discrimination between predictive and non-predictive threat stimuli decrease as threat intensity increases. The central mechanisms that mediate the transition from discriminatory to generalized threat responding remain poorly resolved. Here, we identify the stress- and dysphoria-associated kappa opioid receptor (KOR) and its ligand dynorphin (Dyn), acting in the ventral tegmental area (VTA), as a key substrate for regulating threat generalization. We identified several dynorphinergic inputs to the VTA and demonstrate that projections from the bed nucleus of the stria terminalis (BNST) and dorsal raphe nucleus (DRN) both contribute to anxiety-like behavior but differentially affect threat generalization. These data demonstrate that conditioned threat discrimination has an inverted ‘U’ relationship with threat intensity and establish a role for KOR/Dyn signaling in the midbrain for promoting threat generalization.

## Introduction

In Pavlovian threat conditioning an unconditioned stimulus (US), typically a foot shock, is paired with a conditioned stimulus (CS+), usually an auditory, visual, or olfactory cue [1, 2]. A perceptually distinct unpaired sensory stimulus (CS−) can be interleaved with CS+/US pairings to assess discriminatory learning and memory [2]. In discriminatory threat conditioning as US intensity increases the conditioned response (CR), measured as freezing behavior, also increases and the discrimination between the CS+ and CS− decreases [3–6]. This impairment in discriminatory responding is proposed as a behavioral correlate of generalized fear and anxiety [7].

The dopamine neurotransmitter system has emerged as an important modulator of fear-related learning [8, 9] and threat discrimination [6, 10–12]. We proposed that threat generalization is mediated, in part, by increased uncertainty caused by the dysphoric effects of stress associated with increasing US intensity and subsequent diminution of discriminatory threat coding by VTA dopamine neurons [6, 13]. KOR/Dyn signaling is a key mediator of the dysphoric effects of stress [14] that involves KOR signaling in the VTA [15]. Thus, we hypothesized that KOR/Dyn signaling in the VTA is an important contributor to impaired conditioned threat discrimination at high US threat intensity. Consistent with this hypothesis, we report here that inactivation of KOR signaling through mutagenesis of the KOR gene *Oprk1* in dopamine neurons, or inactivation of the gene encoding preprodynorphin, *Pdyn*, in inputs to the VTA, attenuates threat generalization at high US intensity. We further demonstrate that *Pdyn*-expressing inputs to the VTA from the DRN, but not the BNST are sufficient to impair threat discrimination at lower US intensities. These findings have broad implications for the neural mechanisms of threat discrimination and generalized fear.

## Results

### KOR activation helps drive impairments in threat discrimination

To establish whether conditioned threat discrimination is regulated by KOR signaling, we conditioned independent cohorts of mice with US intensities ranging from 0.1-0.5 mA. One hour prior to the start of the conditioning paradigm, mice were injected with the long-acting KOR antagonist norBNI (10 mg/kg) or saline. Following baseline freezing analysis in context A, mice were conditioned for two days with 10-CS/US pairings in context B and tested 24 hrs later in context A (Figure 1A). Pre-injection with norBNI did not alter US responsiveness (Figure S1) or freezing during conditioning (Figure S2). During testing we observed no difference between groups in response to the CS+ (Figure 1B), but we did observe a significant interaction between US intensity and CS− freezing (Figure 1C). Mice pre-treated with norBNI had significantly less freezing to the CS− relative to saline injected controls at both 0.4 and 0.5 mA US intensities. Analysis of discriminatory threat responses (CS+ minus CS−) revealed a curvilinear or inverted ‘U’ relationship with US intensity that was significantly attenuated at high US intensity by norBNI (Figure 1D).

**Figure 1.**
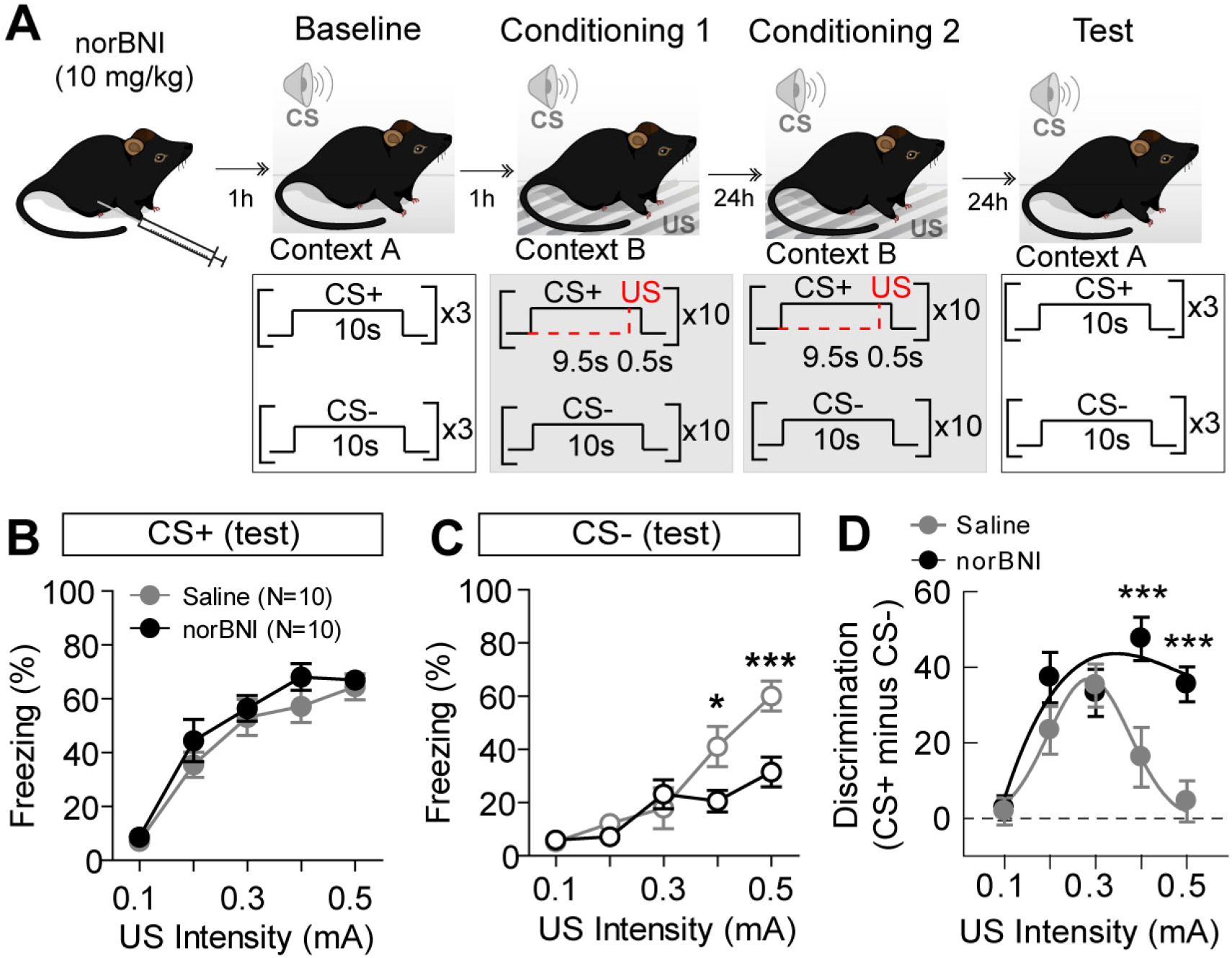
KOR modulation of threat discrimination. (A) Schematic of experimental paradigm. (B) Freezing to CS+ during test in context A. (C) Freezing to CS− during test in context A (N=10 mice per group; Two-way ANOVA, F_*(4,90)*_=4.15, P=0.0039; Bonferroni’s multiple comparisons, *P<0.05, ***P<0.001). (D) Discrimination index (CS+ minus CS−) during test (Two-way ANOVA, F_*(4,90)*_=4.31, P=0.0031; Bonferroni’s multiple comparisons, ***P<0.001). Data are presented as mean ± standard error of the mean (S.E.M).

### KOR signaling in dopamine neurons impairs threat discrimination

Dopamine plays a key role in discriminatory threat conditioning [11] that is sensitive to US intensity [6]. KOR potently modulates the activity of midbrain dopamine neurons [16–19]; thus, we hypothesized that KOR signaling in dopamine neurons contributes to conditioned threat generalization at high US intensity. To test this, we selectively mutated *Oprk1* in dopamine-producing neurons using a single adeno-associated viral vector (AAV) CRISPR/SaCas9 system for conditional expression of SaCas9 and a single guide RNA (sgRNA) [20]. Mice expressing Cre-recombinase under control of the endogenous dopamine transporter (DAT) locus (*Slc6a3*-Cre, or DAT-Cre) were injected with AAV1-FLEX-SaCas9-U6-sg*Oprk1*, or an sgRNA control, and an AAV containing an expression cassette for the nuclear envelope localized EGFP [21] (AAV-FLEX-EGFP-KASH) into the VTA (Figure 2A-B). Four weeks following injection, EGFP-KASH labeled nuclei were isolated and subjected to fluorescence activated cell sorting (FACS). Isolated whole genomic DNA was amplified (WGA) and targeted deep sequencing of a PCR amplicon containing the targeted region was analyzed (Figure 2C). SaCas9 induced a variety of indels in just over 80% of sequence reads (Figure 2C-E). GFP-negative nuclei and predicted off-target sites contained virtually no detectable indels (Figure S3).

**Figure 2.**
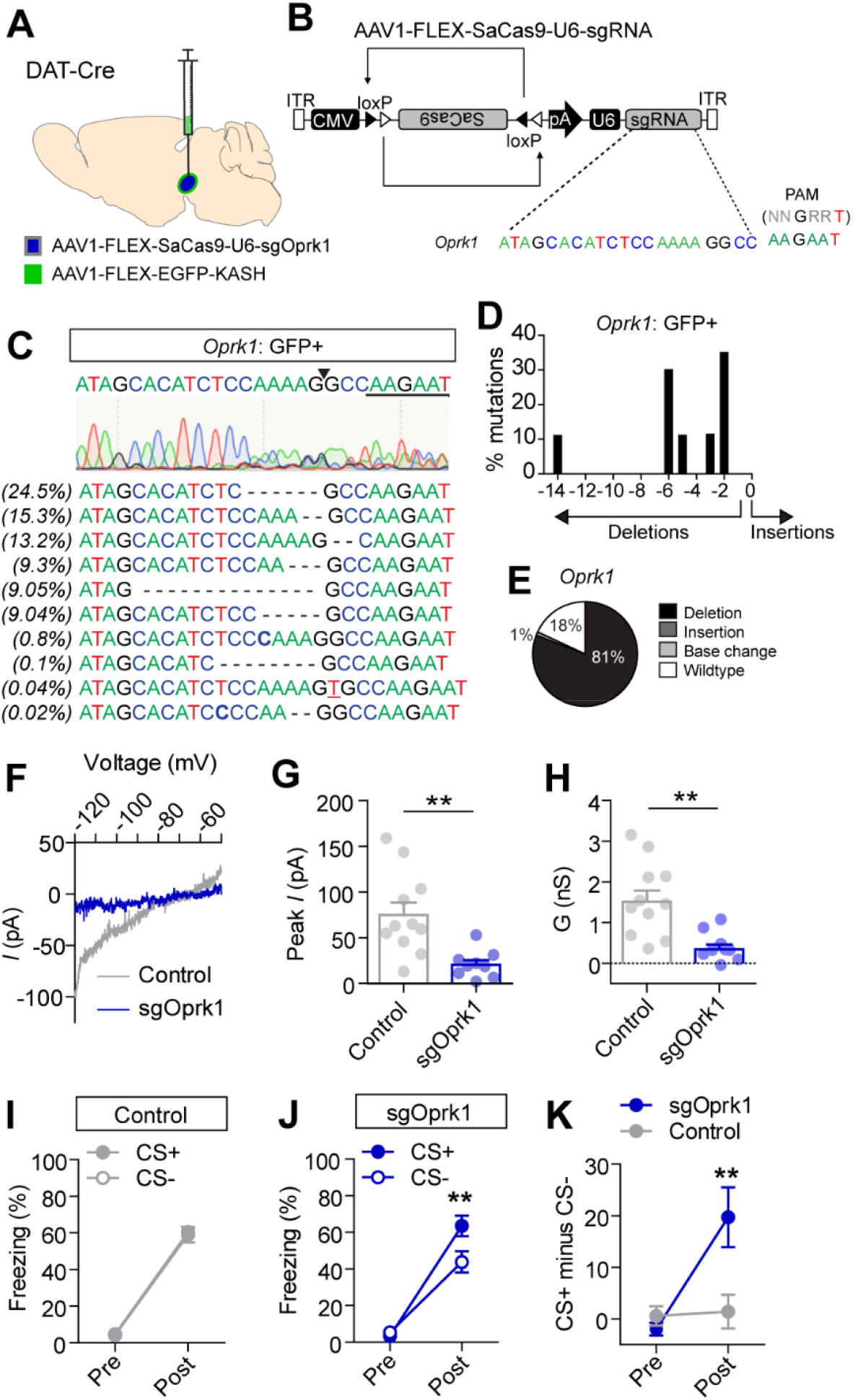
Mutagenesis of *Oprk1* in dopamine neurons. (A-B) Schematic of viral injection location (A) and AAV1-FLEX-SaCas9-U6-sgRNA containing guide for *Oprk1* (B). (C) *Oprk1* guide sequence and predicted cut site (arrowhead) with corresponding sanger sequencing of PCR amplicon (top). Bottom: sequence reads from targeted deep sequencing with percentage of occurrence on left. (D) Distribution of indel mutations from targeted deep sequencing. (E) Proportion of sequence reads with insertions, deletions, base changes, or wild type sequence. (F) Representative subtracted current (*I*)-voltage (*V*) plot (ACSF-U69,593) in a cell from *Oprk1*-targeted or control mouse. (G-H) Peak subtracted *I* (G; Two-tailed Student’s t-test, t_(*18*)_=3.47, P=0.0028) and conductance (H) in cells from control and sg*Oprk1* mice (H; Two-tailed Student’s t-test, t_(*18*)_=3.63, P=0.0019; n=11 cells from N=3 control and n=9 cells from N=3 sg*Oprk1* mice). (I) Freezing to CS+ and CS− during baseline (pre) and test (post) in control mice (N=9) after conditioning with 0.5mA foot shock. (J) Freezing to CS+ and CS− pre- and post-conditioning in sg*Oprk1* mice (N=12; Two-way repeated measures ANOVA, F_(*1,22*)_=7.54, P=0.012; Bonferonni’s multiple comparisons **P<0.01). (K) Discrimination index pre-and post-conditioning in control and sg*Oprk1* mice (Two-way repeated measures ANOVA, F_(*1,19*)_=8.22, P=0.0099; Bonferonni’s multiple comparisons **P<0.01). Data are presented as mean ± S.E.M.

Consistent with the high rate of mutagenesis, whole-cell patch clamp recordings of hyperpolarizing current in dopamine neurons in response to the KOR agonist U69,593 (1 μM) were significantly reduced in *Oprk1* mutant relative to control mice (Figure 2F-H). To assess the impact of *Oprk1* mutagenesis in dopamine neurons, mice were conditioned using a 0.5 mA foot shock as described above (Figure 1a). *Oprk1* inactivation significantly improved CS+/CS− discrimination (Figure 2I-K) that was associated with a reduction in CS− freezing relative to controls (Figure S3). *Oprk1* mutagenesis did not alter US responsiveness, pain sensitivity, conditioned threat acquisition, or anxiety (Figure S3).

### Pdyn inputs to the VTA drive impairments in threat discrimination

KOR signaling is localized to both dopamine neuron terminals and cell bodies [15–19, 22]. Previous analysis of dopamine neuron activity during conditioned threat generalization revealed a reduction in phasic activation in response to cues associated with high US intensity [6]. Therefore, we hypothesized that KOR-dependent suppression of threat discrimination is mediated by Dyn release into the VTA. To establish the location of Dyn inputs to the VTA, we injected the retrograde transducing virus CAV2 containing a conditional expression cassette for the bright fluorescent protein ZsGreen (CAV2-FLEX-ZsGreen; [23]) into the VTA of *Pdyn-*Cre mice (Figure 3A). We identified three sources of Dyn, the nucleus accumbens (NAc), BNST, and the DRN (Figure 3B). *Pdyn-*expressing inputs from the DRN were significantly higher than the other two brain regions (Figure 3C). ZsGreen-positive neurons within the NAc were sparse and broadly distributed. In contrast, inputs from both the BNST and DRN were localized to specific subdivisions of these structures (Figure 3D-E).

**Figure 3.**
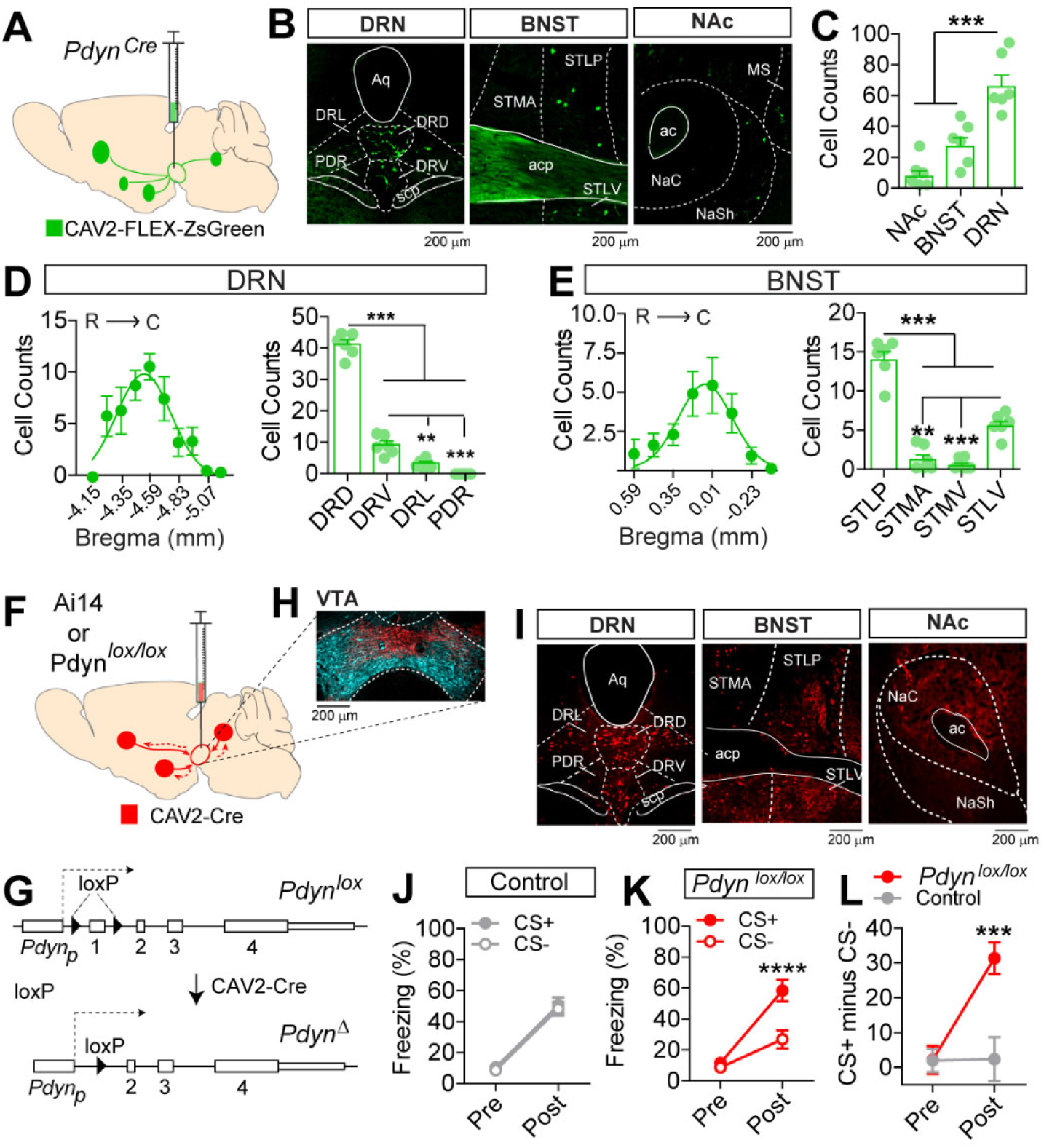
Inactivation of *Pdyn* in the inputs to the VTA. (A) Schematic of CAV2-FLEX-ZsGreen injection into *Pdyn*^*Cre*^ mice. (B) Representative images of ZsGreen fluorescence in the DRN, BNST, and NAc. (C) Normalized cell counts of DRN-*Pdyn* inputs are significantly higher than NAc or BNST inputs (N=6 mice; One-way ANOVA, F_(*2,15*)_=25.01, P<0.0001, Tukey’s multiple comparisons, ****P<0.0001). (D) Normalized distribution of *Pdyn*-expressing neurons in subregions of the DRN (One-way ANOVA, F_(*3,20*)_=347.07, P<0.0001, Tukey’s multiple comparisons, ***P<0.0001, **P<0.01). (E) Normalized distribution of *Pdyn*-expressing neurons in subregions the BNST (One-way ANOVA, F_(*3,20*)_=78.57, P<0.0001, Tukey’s multiple comparisons, ***P<0.0001, **P<0.01). (F) Schematic of CAV2-Cre injection into the VTA of *Pdyn*^*lox/lox*^ or Ai14 mice. Schematic of *Pdyn*^*lox*^ allele before and after Cre-mediated recombination. (H) Representative image of TdTomato (red) and TH (cyan) fluorescence in VTA of Ai14 mouse injected with CAV2-Cre. (I) Representative image of TdTomato fluorescence in DRN, BNST, and NAc of Ai14 mouse injected with CAV2-Cre. (J) Freezing to CS+ and CS− during baseline (pre) and test (post) in control mice (N=10). (K) Freezing to CS+ and CS− pre- and post-conditioning (0.5 mA foot shock) in CAV2-Cre::*Pdyn*^*lox/lox*^ mice (N=12; Two-way repeated measures ANOVA, F_(*1,22*)_=12.55, P=0.0018; Bonferonni’s multiple comparisons ****P<0.0001). (L) Discrimination index pre-and post-conditioning in control and CAV2-Cre::*Pdyn*^*lox/lox*^ mice (Two-way repeated measures ANOVA, F_(*1,20*)_=13.15, P=0.0017; Bonferonni’s multiple comparisons ***P<0.01). Data are presented as mean ± S.E.M. Abbreviations: DRD: dorsal raphe dorsal, DRV: dorsal raphe ventral, DRL: dorsal raphe lateral, PDR: posterior dorsal raphe, Aq: cerebral aqueduct, scp: superior cerebellar peduncle, STLP: stria terminalis posterior, STLV: stria terminalis ventral, STMA: stria terminalis medial anterior, STMV: stria terminalis medial ventral, ac: anterior commissure, acp: anterior commissure posterior.

To inactivate *Pdyn* in inputs to the VTA, *Pdyn*^*lox/lox*^ mice [24] were injected with CAV2-Cre [25] into the VTA (Figure 3F-G). CAV2-mediated recombination occurred in the NAc, BNST, and DRN as illustrated in the TdTomato reporter line Ai14 [26] (Figure 3H-I). Similar to mutagenesis of *Oprk1* in dopamine neurons, inactivation of *Pdyn* in inputs to the VTA significantly improved conditioned threat discrimination in mice conditioned with a US intensity of 0.5 mA (Figure 3J-L). Inactivation of *Pdyn* did not alter US responsiveness, pain sensitivity, acquisition of conditioned threat responses, or anxiety (Figure S3). Improved threat discrimination was associated with a significant reduction in freezing to the CS− in CAV2-Cre::*Pdyn*^*lox/lox*^ mice relative to controls (Figure S3).

### Differential effects of Pdyn inputs to the VTA in threat discrimination

To establish whether activation of *Pdyn*-expressing inputs to the VTA are sufficient to impair conditioned threat discrimination, we targeted expression of channel-rhodopsin (ChR2; [27]) to the BNST and DRN. *Pdyn*-Cre mice were injected with AAV1-FLEX-ChR2-mCherry into the BNST or DRN and bilaterally implanted with optical fibers above the VTA (Figure 4A-B). Mice were conditioned for two-consecutive days with 0.3 mA foot shocks and inputs were activated with blue light stimulation at 20 Hz for 1 s beginning 0.5 s prior to US delivery (Figure 4C). Activation of BNST inputs did not affect US responses during conditioning (Figure 4D-E) and was not associated with a change in pain sensitivity (Figure S4). Acquisition of conditioned freezing was also not affected by stimulation of BNST inputs during US presentation (Figure S4). Both control and BNST-*Pdyn*::ChR2 mice had significant discrimination of CS+ and CS− stimuli following conditioning (Figure 4f-h); however, stimulation of BNST inputs increased overall freezing to both the CS+ and CS− (Figure S4). Although stimulation of BNST-*Pdyn* inputs did not impair discrimination, activating these inputs (20 Hz stimulation, 3s on 3s off) during exploration of an elevated-plus maze (EPM) was associated with an increase in anxiety-like behavior as measured by time in the open arm (Figure 4I). Reduced time in the open arm of the EPM corresponded with a slight, but significant reduction in overall distance traveled during the assay (Figure S4).

**Figure 4.**
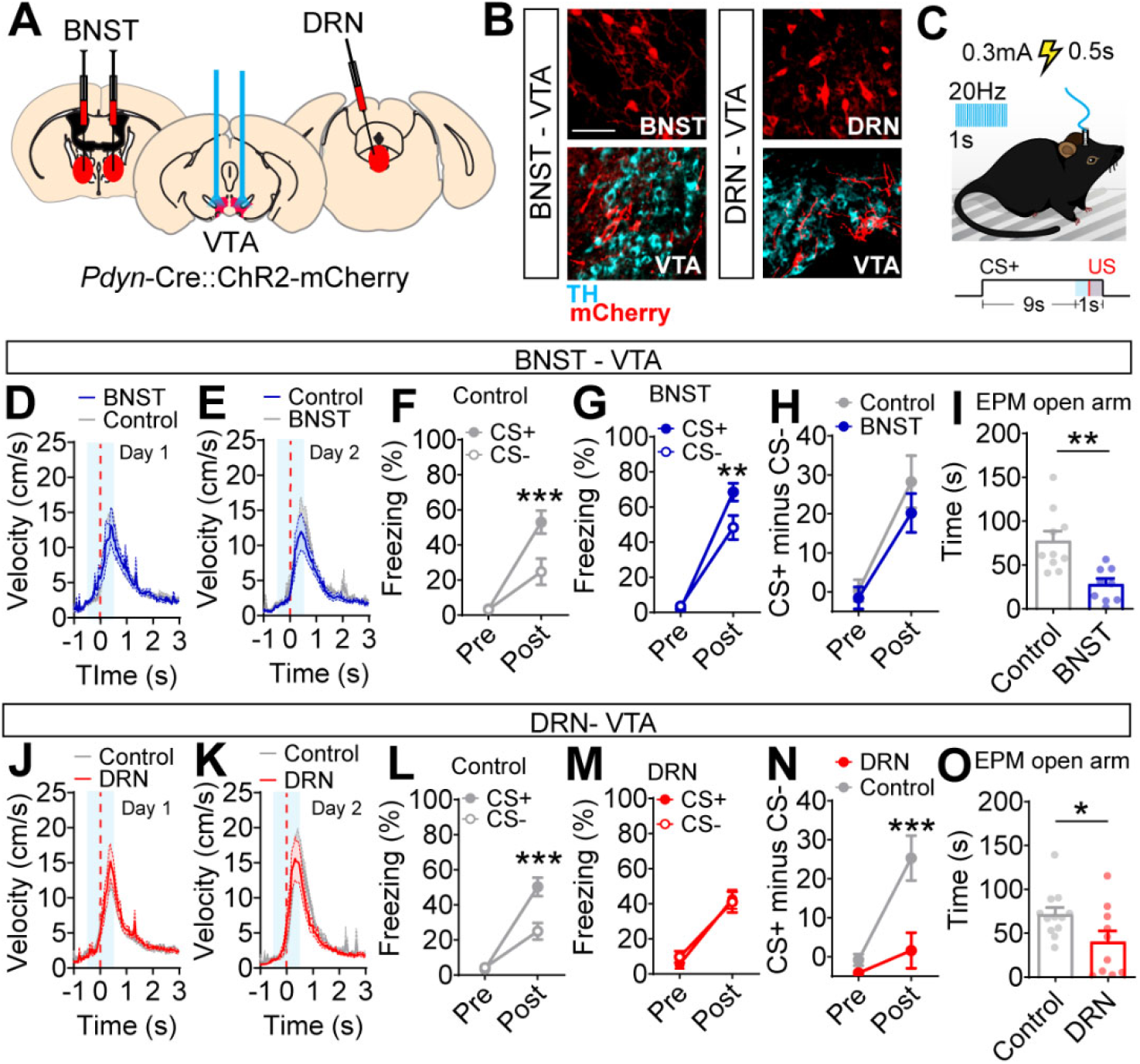
Stimulation of *Pdyn* inputs to the VTA. (A) Schematic illustration of AAV1-FLEX-ChR2-mCherry injection into the BNST or DRN and bilateral optical cannula implantation into the VTA. (B) Representative histological verification of ChR2-mCherry expression in the BNST or DRN and ChR2-mCherry fibers in the VTA; scale bar = 50 μm. (C) Schematic illustration of experimental paradigm with 20 Hz optical stimulation (474 nm, 0.5 ms pulse width) for 1 s beginning 0.5 s before footshock (0.3 mA) onset and co-terminating with US and CS+. (D-E) Velocity of movement during foot shock presentation on day 1 (D) and day 2 (E); note control and BNST stimulation curves are nearly completely superimposed. (F) Pre- and post-conditioning freezing in BNST-*Pdyn*::mCherry control mice (N=10; Two-way repeated measures ANOVA, F_(*1,18*)_=9.36, P=0.0067, Bonferonni’s multiple comparisons ***P<0.001). (G) Pre- and post-conditioning freezing in BNST-*Pdyn*::ChR2-mCherry mice (N=9; Two-way repeated measures ANOVA, F_(*1,16*)_=6.35, P=0.023, Bonferonni’s multiple comparisons **P<0.01). Discrimination index pre- and post-conditioning. (I) Time spent in open arm of EPM (N=10 control and N=9 BNST, two-tailed, unpaired Student’s t-test, t_(17)_=3.75, **P=0.0016). (J-K) Velocity of movement during foot shock presentation (0.3 mA) and stimulation on day 1 (J) and day 2 (K); note control and DRN stimulation curves are nearly completely superimposed. (L) Pre- and post-conditioning freezing in DRN-*Pdyn*::mCherry control mice (N=12; Two-way repeated measures ANOVA, F_(*1,22*)_=12.99, P=0.0016, Bonferonni’s multiple comparisons ***P<0.001). (M) Pre- and post-conditioning freezing in DRN-*Pdyn*::ChR2-mCherry mice (N=10). (N) Discrimination index pre- and post-conditioning (Two-way repeated measures ANOVA, F_(*1,20*)_=5.78, P=0.026, Bonferonni’s multiple comparisons ***P<0.001). (O) Time spent in open arm of EPM (N=12 control and N=10 DRN, two-tailed, unpaired Student’s t-test, t_(20)_=2.20, *P=0.04). Data are presented as mean ± S.E.M.

Stimulation of DRN-*Pdyn* inputs to the VTA did not affect US responses during conditioning (Figure 4J-K) and was not associated with a change in pain sensitivity (Figure S4), similar to stimulation of BNST-*Pdyn* inputs. Acquisition of conditioned freezing was also not affected by stimulation of DRN inputs during US presentation (Figure S4). Optogenetic stimulation of DRN-*Pdyn* inputs during conditioning impaired threat discrimination in the post conditioning test (Figure 4L-N) that was associated with a significant increase in freezing to the CS− without affecting freezing to the CS+ (Figure S4). Like BNST inputs, stimulation of DRN-*Pdyn* inputs increased anxiety-like behavior and reduced exploration in the EPM test (Figure 4O and FigureS4).

## Discussion

Our data demonstrate that conditioned threat discrimination follows an inverted ‘U’ relationship with US intensity. These results are consistent with the Yerkes-Dodson law [28], or at least variations in the interpretation of this law as it pertains to increasing stress, arousal, or motivation [29]. Our results further support the role of KOR signaling in the dysphoric effects of stress [14] at the level of the VTA [15] for promoting generalization as threat intensity increases.

The dysphoric effects of KOR/Dyn signaling is proposed to contribute to anxiety-like behavior [30]. We find that stimulation of both BNST and DRN inputs to the VTA increase anxiety-like behavior, but only activation of DRN inputs to the VTA are sufficient to disrupt threat discrimination. This suggests an important dissociation between threat generalization and anxiety and point to potentially shared circuitry for *PDyn*:DRN→VTA and *Pdyn*:BNST→VTA input/output relationships for anxiety that diverge at the level of regulation of threat discrimination. These inputs may also differentially regulate outputs based on previously described differences in projection-specific somatodendritic modulation of VTA dopamine neurons activity [17, 18].

In addition to our observations that *PDyn*:DRN→VTA and *Pdyn*:BNST→VTA inputs regulate anxiety, KOR/Dyn signaling in the DRN and BNST has also been linked to stress and anxiety [31–34].Thus, KOR signaling is likely key to the coordinated actions of the BNST, DRN, and VTA. The nature of KOR modulation of threat discrimination coding by the VTA remains to be established. Given the inhibitory nature of KOR signaling at the level of the VTA [15–19] it is likely that increasing Dyn release into the VTA in response to increasing US intensity diminishes the capacity of dopamine neurons to provide a modulatory signal for threat discrimination learning [6].

## Methods

### Mice

All experiments were approved and in accordance with the guidelines of the Institutional Animal Care and Use Committee of the University of Washington. Similar numbers of male and female mice of 8 weeks or older were used for all behavioral experiments. Mice were group housed (2 - 5 mice per cage), and kept on a 12-hour light/dark cycle (lights on at 7 am). Access to food and water was *ad libitum*. All experiments were performed during the light cycle and replicated in at least 2 cohorts. All mice were bred on a C57BL/6J background. *Pdyn*^*IRES-Cre*^ mice and *Pdyn*^*lox/lox*^ mice were provided by Dr. Richard Palmiter, University of Washington. *Slc6a3*^*IRES-Cre*^ (DAT-Cre, stock# 006660) and Ai14 (*Gt(ROSA)26Sor*^*tm14(CAG-tdTomato)Hze*^, stock#007908) were obtained from Jackson Laboratories.

### Viral production

All CAV and AAV viruses were produced in house with titers of 1-3 ×10^12^ particles per mL as described [35].

### Surgery

Mice were anesthetized with isoflurane and positioned on a stereotaxic alignment device (David Kopf Instruments). During the surgery, 1.5-2.0% isoflurane was continuously administered through a nose port and body temperature was maintained with a heating pad. All anterior-posterior coordinates were adjusted to the distance between Bregma and Lambda (A-P coordinate * (Bregma-Lambda distance/4.21)). After small burr holes were drilled, the injection needle was lowered straight down or at an angle (for DRN), the needle was left for 2 minutes and then pulled up 0.4 mm while virus was injected at a rate of 0.25 μl/min. The bilateral viral injection coordinates from Bregma in millimeters were: VTA, A-P: −3.25; M-L: ±0.5, D-V: −4.5; DRN, A-P: −4.55, M-L: −0.3 at a 5° angle, D-V: −3.2; BNST, A-P: +0.2, M-L: ±1.250, D-V: −4.25 & A-P: −0.15, M-L: ±1.250, D-V: −4.00. For *in vivo* ChR_2_ experiments, optic fibers were placed 0.6 mm above the D-V coordinates. For VTA bilateral optic fiber placement, one optic fiber was inserted at a 10° angle, and the M-L coordinate was adjusted to −1.225, and the respective D-V coordinate to −4.060. All optic fibers were fabricated in house [36] and fixed to the skull with dental cement. Only those fibers that had a light loss not exceeding 30% were used. Prior to behavioral testing, mice were allowed to recover for at least two weeks.

### Retrograde Input Mapping

Three weeks following injection of 500 nl CAV2-FLEX-zsGreen into the VTA of *Pdyn*^*IRES-Cre*^ mice or CAV2-Cre into the VTA of Ai14 mice, animals were euthanized and perfused with 4% paraformaldehyde. 30 μm frozen brain sections were collected, mounted on glass slides, and coverslipped. Brain sections corresponding to a reference atlas (approximately every third section) were imaged at 10x magnification using a Keyence BZ-X710 fluorescent microscope. Cells were counted bilaterally for each section corresponding to areas where fluorescent cells were located. Cell counts were normalized to the average total number of cells counted per animal.

### NorBNI treatment

For drug injection experiments, animals were intraperitoneal (i.p.) injected (0.1 ml/ 10 g) 1 hour prior to baseline freezing assessment with norbinalthorphimine (norBNI, Tocris Cat.# 0347) diluted in sterile saline (1 mg/ ml) or saline (control).

### Behavioral Assays

The start time of assays were kept consistent for each cohort. Animals were placed in the testing room at least 1 hour before the starting time of the sessions for habituation. All mice were handled prior to testing and, if relevant, to the attachment and detachment of the patch cord for at least 3 days prior to fear conditioning. For *in vivo* ChR2 stimulation during fear conditioning, the blue light parameters were as follows: 20 Hz, 5 ms pulse-width, for 21 pulses starting 500 ms before US onset. For all other *in vivo* ChR2 behavioral experiments, the blue light stimulation was 20 Hz, 10 ms pulse-width, 3 seconds on, 3 seconds off.

#### Fear conditioning

Identical fear conditioning chambers (21.6 × 17.8 × 12.7 cm; Med Associates Inc.) were situated inside sound-attenuating boxes. Each chamber had three aluminum and one Plexiglas wall. The speaker for auditory tones was placed on one aluminum wall. The floor consisted of 24 stainless steel rods connected to a scrambled shock generator. A video camera was mounted to the ceiling of the sound-attenuating box to record animal behavior and movement. The chambers were cleaned using 70% ethanol between animal conditioning sessions. For the baseline measurements and fear retention a white cardboard insert was placed inside the chamber and the box was cleaned with 1% acetic acid between subjects to prevent contextual learning. For the fear conditioning paradigm, a baseline freezing measurement session was first conducted. During this session, animals were introduced to the counterbalanced assigned CS+ and CS− tones (4 and 12 kHz, 10 secs). After a 2-minute habituation period, the tones were presented three times each in a serial order (CS+ was always presented on the first trial; ITI = 60 secs). 60 seconds after the last tone presentation, the mice were returned to their home cages. After 2 hours, the mice were placed in the conditioning chamber (aluminum walls). Conditioning also began with a 2-minute habituation period. The CS+ (10 secs; ITI = 60 secs) was always presented on the first trial and paired with a 0.5- s foot shock of 0.1, 0.2, 0.3, 0.4, or 0.5 mA. The CS− (10 secs) was never paired with a foot shock. Each cue was presented 10 times during conditioning. On the consecutive day after final conditioning, a retention session (same as baseline measurement session) was performed. Movement velocities (cm/ s) were calculated offline using video tracking software Ethovision XT 8.5 (Noldus Information Technology). Freezing was analyzed using custom MATLAB script and parameters were set at a velocity of less than 0.75 cm/s for the duration of at least 1 s [6]. Freezing behavior before, during and after CS+ and CS− was scored.

#### Elevated plus maze

The elevated plus maze (EPM) was elevated 120 cm above the floor and consisted of two open arms and two black plexiglass closed arms (30 cm length, 6 cm width, 15 cm high walls). Mice were placed in the center zone of the EPM facing one of the open arms and were allowed to explore freely for 5 minutes. For optogenetic experiments, inputs to the VTA were stimulated at 20 Hz, 10 s on, 10 s off. Their behavior was recorded by a camera installed to the ceiling of the experimental room. The time spend (s) and distance travelled (cm) in each zone (open arms, closed arms and center) was calculated offline with video tracking software Ethovision XT 8.5 (Noldus Information Technology).

#### Hot plate

The hot plate was set at a constant temperature of 57.5 °C. The time spend (cs) on the hotplate was recorded manually from the moment upon which the 4 paws touched the hotplate until the animal jumped or licked its paws. For optogenetic experiments, inputs to the VTA were stimulated at 20 Hz, 10 s on, 10 s off. The mouse was removed as soon as the timer was stopped.

### Immunohistochemistry

Mice were deeply anaesthetized with 5 mg/kg Beuthanasia-D (Merck Sharp and Dohme Corp) and transcardially perfused with cold 1 x phosphate buffered saline (PBS) followed by cold 4% paraformaldehyde (PFA) in 1 x PBS. Brains were immediately extracted, post fixed in 4% PFA at 4°C overnight and then transferred and stored in 30% sucrose in 1 × PBS at 4°C until further use. Brains were sectioned with a cryostat (Leica CM 1850) in 30μm sections at −20 °C and stored free-floating in 0.02% Sodium Azide in 1 X PBS. For immunohistochemistry, slices were blocked in 0.3% Triton and 3% Normal Donkey Serum (NDS) in 1 x TBS, and then incubated in primary antibody anti-ZsGreen (rabbit, 1:2000, Invitrogen A11122) at 4°C overnight. The slices were then washed with 1 x PBS three times and incubated for 1 hour at room temperature with a secondary antibody conjugated to AlexaFluor 488 (anti-rabbit, 1:200, Jackson Immunolabs). The sections were washed in 1 x PBS three times and mounted and then cover-slipped with Fluoromount-G with DAPI (Southern Biotech).

### Generation and validation of AAV1-FLEX-SaCas9-U6-sgOprk1

The sgRNA targeted to the *Oprk1* locus (sg*Oprk1*) was designed as previously described [20]. The following oligos (Sigma) were used to clone into pAAV-FLEX-SaCas9-U6-sgRNA (Addgene 124844). *Oprk1* forward: CACCGATAGCACATCTCCAAAAGGCC; *Oprk1* reverse: AAACGGCCTTTTGGAGATGTGCTATC.

#### Targeted deep sequencing of Oprk1 locus

AAV1-FLEX-SaCas9-sg*Oprk1* and AAV1-FLEX-EGFP-KASH were co-injected into the VTA of DAT-Cre mice. Nuclei isolation, FACS, and targeted deep sequencing were performed as described previously [20]. Four weeks following surgery, tissue punches of the ventral midbrain from 3 mice were pooled into a single group and homogenized in 2mL of homogenization buffer containing (in mM): 320 Sucrose (sterile filtered), 5 CaCl (sterile filtered), 3 Mg(Ac)2 (sterile filtered), 10 Tris pH 7.8 (sterile filtered), 0.1 EDTA pH 8 (sterile filtered), 0.1% NP40, 0.1 Protease Inhibitor Cocktail (PIC, Sigma), 1 β-mercaptoethanol. The volume of the homogenate was brought up to 5mL using homogenization buffer, mixed by inversion, and incubated on ice for 5 minutes. 5mL of 50% Optiprep density gradient medium (Sigma) containing (in mM): 5 CaCl (sterile filtered), 3 Mg(Ac)2 (sterile filtered), 10 Tris pH 7.8 (sterile filtered), 0.1 PIC, 1 β-mercaptoethanol was added to the homogenate and mixed by inversion. The mixture was gently loaded on 10mL of 29% iso-osmolar Optiprep solution in a 1×3 ½ in a Beckman centrifuge tube (SW32 Ti rotor) and spun at 7500 RPM for 30min at 4°C. The floating cell debris was removed using a KimWipe and the supernatant was gently poured out. The nuclei pellet was vigorously resuspended in sterile 1xPBS. GFP-positive nuclei were sorted using a BD AriaFACS III. DNA was extracted and purified using a phenol chloroform extraction. Whole genome amplification (WGA) was performed using the REPLI-g Advanced DNA Single Cell kit (Qiagen) according to manufacturer’s instructions.

For generation of the specific amplicons, 1ul of WGA DNA was diluted 1:50 and amplified (PCR1) with Phusion High Fidelity Polymerase (Thermo Fisher) using the following thermocycler protocol: initial denaturation (30sec, 95°C); denaturation (10sec, 95°C); annealing (20sec, 66°C); extension (10sec, 72°C); cycle repeated x34; final extension (5min, 72°C). For PCR2, 1uL of PCR 1 was amplified with a second set of primers using the same thermocycler protocol. The 240bp amplicon from PCR2 was gel extracted using the MinElute gel extraction kit (Qiagen), and 500ng of product were sent to Genewiz for Amplicon-EZ targeted deep sequencing. The primers used for the generation of the amplicons were as follows: PCR1 forward:

AAAGCACTGAGATAGAACCT reverse: TTCCAAAGTAAAAGTTCACC, PCR2 forward: GTTTAAGTTGCAGACATGG; reverse: TCAAGGTGAATATGCTGG.

### Slice Electrophysiology

DAT-Cre mice at 6 weeks of age were injected in the VTA with AAV-FLEX-YFP and AAV-FLEX-saCas9-U6-sgOprk1 or a control CRISPR virus. 4 weeks following viral injection horizontal VTA slices (200 μm) were prepared in an ice slush solution containing (in mM): 92 NMDG, 2.5 KCl, 1.25 NaH_2_PO_4_, 30 NaHCO_3_, 20 HEPES, 25 glucose, 2 thiouria, 5 Na-ascorbate, 3 Na-pyruvate, 0.5 CaCl_2_, 10 MgSO_4_, pH 7.3-7.4 [37]. Slices recovered for ≤12 minutes in the same solution at 32°C and then were transferred to a room temperature solution including (in mM): 92 NaCl, 2.5 KCl, 1.25 NaH_2_PO_4_, 30 NaHCO_3_, 20 HEPES, 25 glucose, 2 thiouria, 5 Na-ascorbate, 3 Na-pyruvate, 2 CaCl_2_, 2 MgSO_4_. Slices recovered for an additional 45 minutes before recordings were made in ACSF at 32°C continually perfused over slices at a rate of ~2 ml/min and containing (in mM): 126 NaCl, 5.5 KCl, 1.2 NaH2PO4, 1.2 MgCl2 11 D-glucose, 18 NaHCO3, 2.4 CaCl2. All solutions were continually bubbled with O_2_/CO_2_.

Whole-cell recordings of fluorescently labeled dopamine neurons were made using an Axopatch 700B amplifier (Molecular Devices) with filtering at 1 KHz using 4-6 MΩ electrodes. Electrodes were filled with an internal solution containing (in mM): 130 K-gluconate, 10 HEPES, 5 NaCl, 1 EGTA, 5 Mg-ATP, 0.5 Na-GTP, pH 7.3, 280 mOsm. Cells were held at −70 mV and currents were evoked using a 10 sec voltage ramp from −120 mV to −50 mV. 3 sweeps were averaged for each cell. U69,593 (1 μM) was bath applied for 10 minutes, followed by 3 additional ramp recordings, which were averaged. The baseline trace was subtracted from the U69,593 trace for each cell and the peak current at −120 mV was measured along with the slope (conductance) using Clampfit software (Molecular Devices).

### Statistical analyses

All statistical testing for behavioral results was performed in Prism (Graphpad) software. Significance between averages for unpaired data was evaluated with student’s t-tests. One-way ANOVAs were performed for multiple groups with a single variable (cell counts). Two-way repeated measures ANOVAs were used for between- or within subject variables (e.g. treatment and US intensity). Where significant interactions were found in an ANOVA, Bonferroni’s multiple comparisons were used for post-hoc pairwise comparisons (Two-way) or Tukey’s multiple comparisons (One-way). For curve-fitting, the best fit was determined for each group by the highest explained variance, i.e. R^2^. Corrected p-values < 0.05 were considered significant. Data is expressed as mean ± SEM unless indicated otherwise.

## Acknowledgments

We thank members of the Zweifel lab for scientific discussion on the design and implementation of experiments. We also thank Dr. James Allen for assistance in the production of AAV viral vectors, Madison Baird for assistance with histological validation, Dr. Charles Chavkin for nonBNI, and Dr. Richard Palmiter for *Pdyn*^*lox/lox*^ and *Pdyn*^*IRES-Cre*^ mice. This work was funded by the US National Institutes of Health (P50MH10642, R01DA044315, and P30DA048736 L.S.Z; F31MH116549 A.C.H.), Stichting Fundatie van de Vrijvrouwe van Renswoude (L.F.) and dr. Hendrik Muller’s Vaderlandsch Fonds (L.F.).

## Author Contributions

L.F., Y.S.J, M.E.S., A.C.H., and L.S.Z designed experiments, collected, and analyzed data and wrote the manuscript. L.F. and Y.S.J. performed viral injection surgeries and behavioral analysis. A.C.H. generated and validated AAV1-FLEX-SaCas9-U6-sg*Oprk1*. M.E.S. performed slice electrophysiology. Histology and cell counts were performed by M.E.S. and L.S.Z. L.S.Z generated CAV2-FLEX-ZsGreen. L.S.Z. and A.C.H. purified all viral vectors.

