## Supplemental Figures for "A midbrain dynorphin circuit promotes threat generalization"

**Summary:** Supplemental Figures for Fellinger et. al.

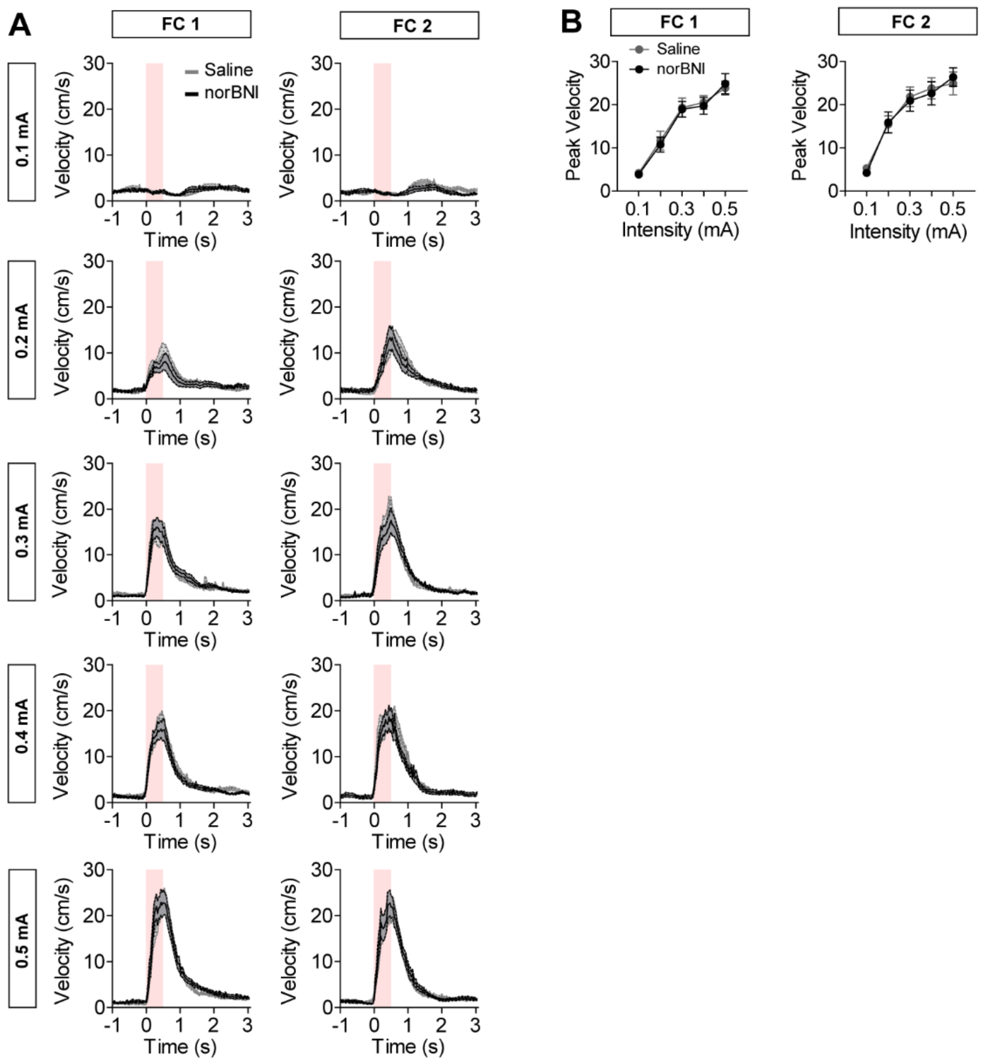


**Figure S1 related to Figure 1. Shock responsiveness in nor-BNI and saline injected mice.** (A) Average velocity of movement during foot shock at different US intensities on days 1 and 2 of fear conditioning (FC). (B) Average peak velocities of mice injected with saline or nor-BNI (A and B, N= 10 mice per group). Data are presented as mean ± S.E.M.

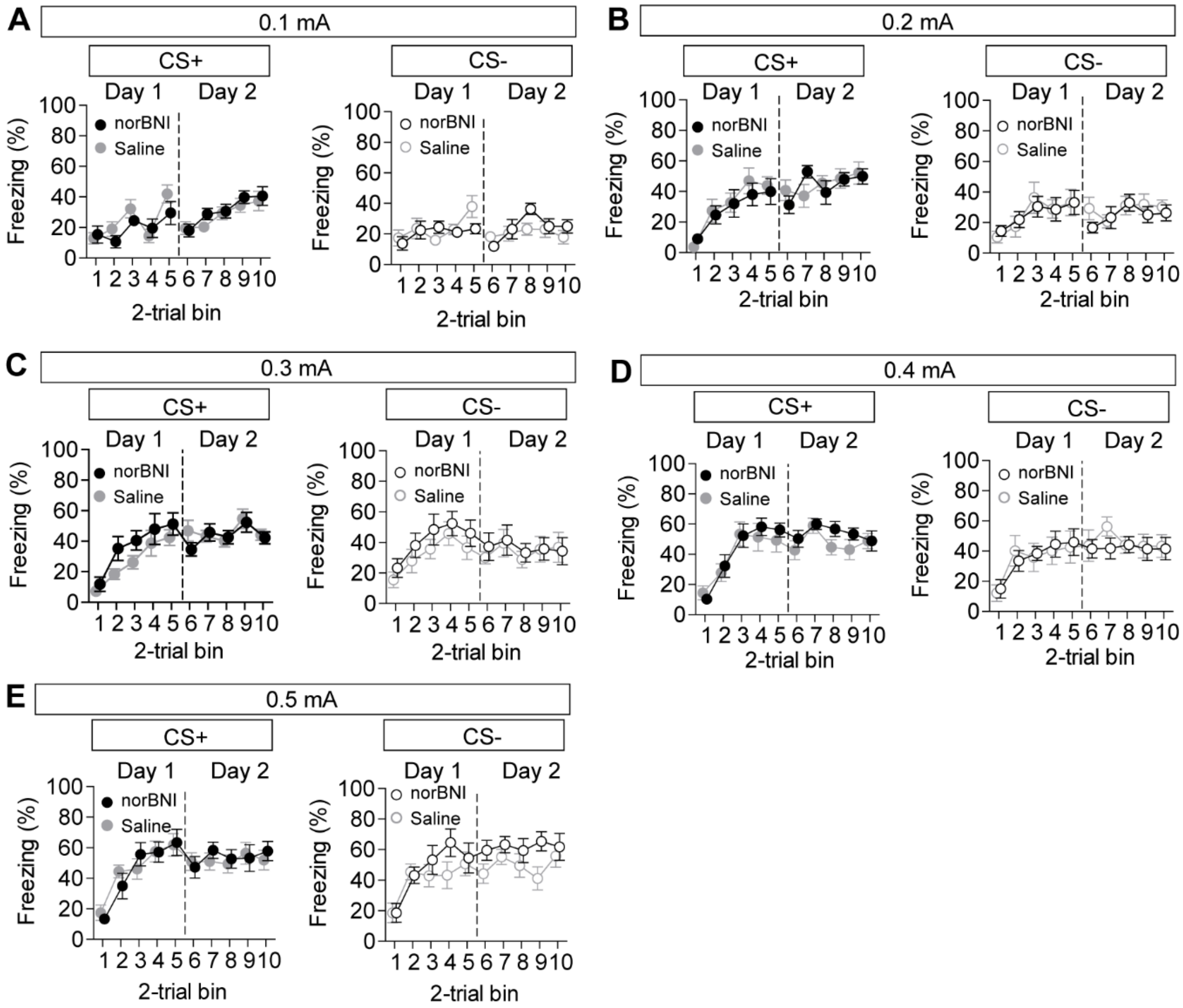


**Figure S2 related to Figure 1. Freezing to CS+ and CS- during conditioning in nor-BNI and saline injected mice.** (A-E) Average freezing behavior during conditioning days 1 and 2 in response to CS+ and CS- in mice injected with saline or nor-BNI at 0.1 (A), 0.2 (B), 0.3 (C), 0.4 (D), and 0.5 mA (E) (N= 10 mice per group). Data are presented as mean ± S.E.M.

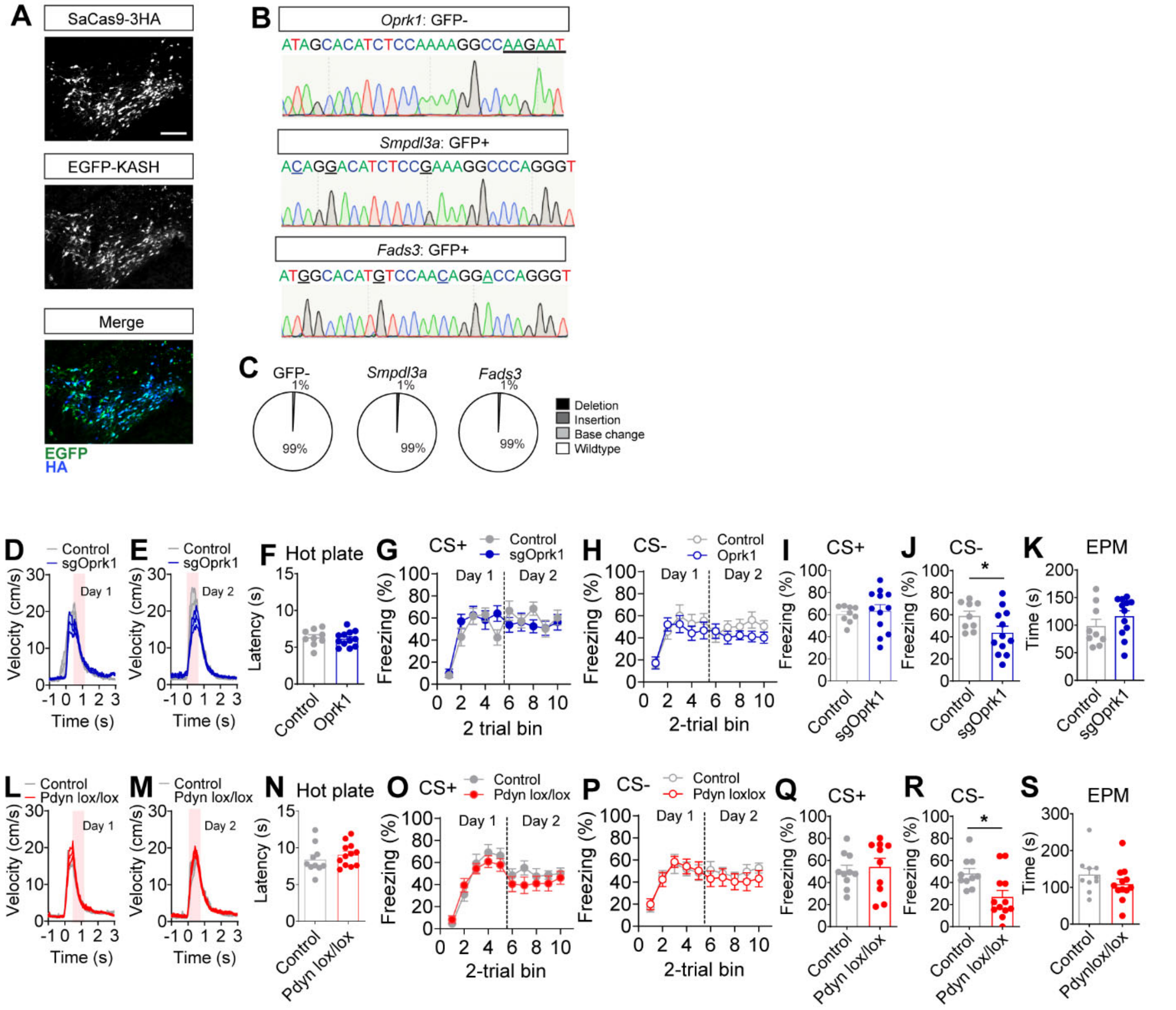


**Figure S3 related to Figure 2. SaCas9 mutagenesis of *Oprk1* and extended behavioral analysis.** (A) Histological verification of HA-tagged SaCas9 and EGFP-KASH expression in the VTA of DAT-Cre mice; scale bar = 100 μm. (B) Sanger sequencing of *Oprk1* in GFP-negative (GFP-) nuclei from FACS of sg*Oprk1*-targeted mice (top) and CRISPOR-predicted off-target genes, *Smpdl3a* and *Fads3*, in GFP-positive (GFP+) neurons from sg*Oprk1*-targeted mice. (C) Proportion of deep sequence reads from PCR amplicons of WGA genomic DNA from GFP- (*Oprk1*) and GFP+ (*Smpdl3a* and *Fads3*) nuclei. (D-E) Average velocity of movement during foot shock at different US intensities on days 1 (D) and 2 (E) of conditioning in sg*Oprk1*-targeted or control DAT-Cre mice. (F) Latency to paw-lick or jump on a 57.5$^0$C hotplate in sg*Oprk1*-targeted or control DAT-Cre mice. (G-H) Average freezing behavior during conditioning days 1 and 2 in response to CS+ (G) and CS- (H) in sg*Oprk1*-targeted or control DAT-Cre mice. (I-J) Freezing to CS+ (I) or CS- (J) on test day in sg*Oprk1*-targeted versus control DAT-Cre mice (J, Two-tailed Student's t-test, $t_{(18)}$=2.12, P=0.049). (K) Time spent in the open arm of an elevated plus maze in sg*Oprk1*-targeted versus control

DAT-Cre mice. (D-K) N=9 control and N=12 sg*Oprk1* mice. (L-M) Average velocity of movement during foot shock at different US intensities on days 1 (L) and 2 (M) of conditioning in CAV2-Cre:*Pdyn*$^{lox/lox}$ or control mice. (N) Latency to paw-lick or jump on a 57.5$^0$C hotplate in CAV2-Cre:*Pdyn*$^{lox/lox}$ or control mice. (O-P) Average freezing behavior during conditioning days 1 and 2 in response to CS+ (O) and CS- (P) in CAV2-Cre:*Pdyn*$^{lox/lox}$ or control mice. (Q-R) Freezing to CS+ (Q) or CS- (R) on test day in CAV2-Cre:*Pdyn*$^{lox/lox}$ or control mice. (R, Two-tailed Student's t-test, $t_{(20)}$=2.78, P=0.012). (S) Time spent in the open arm of an elevated plus maze in CAV2-Cre:*Pdyn*$^{lox/lox}$ or control mice. (L-S) N=10 control and N=12 CAV2-Cre:*Pdyn*$^{lox/lox}$ mice. Data are presented as mean ± S.E.M.

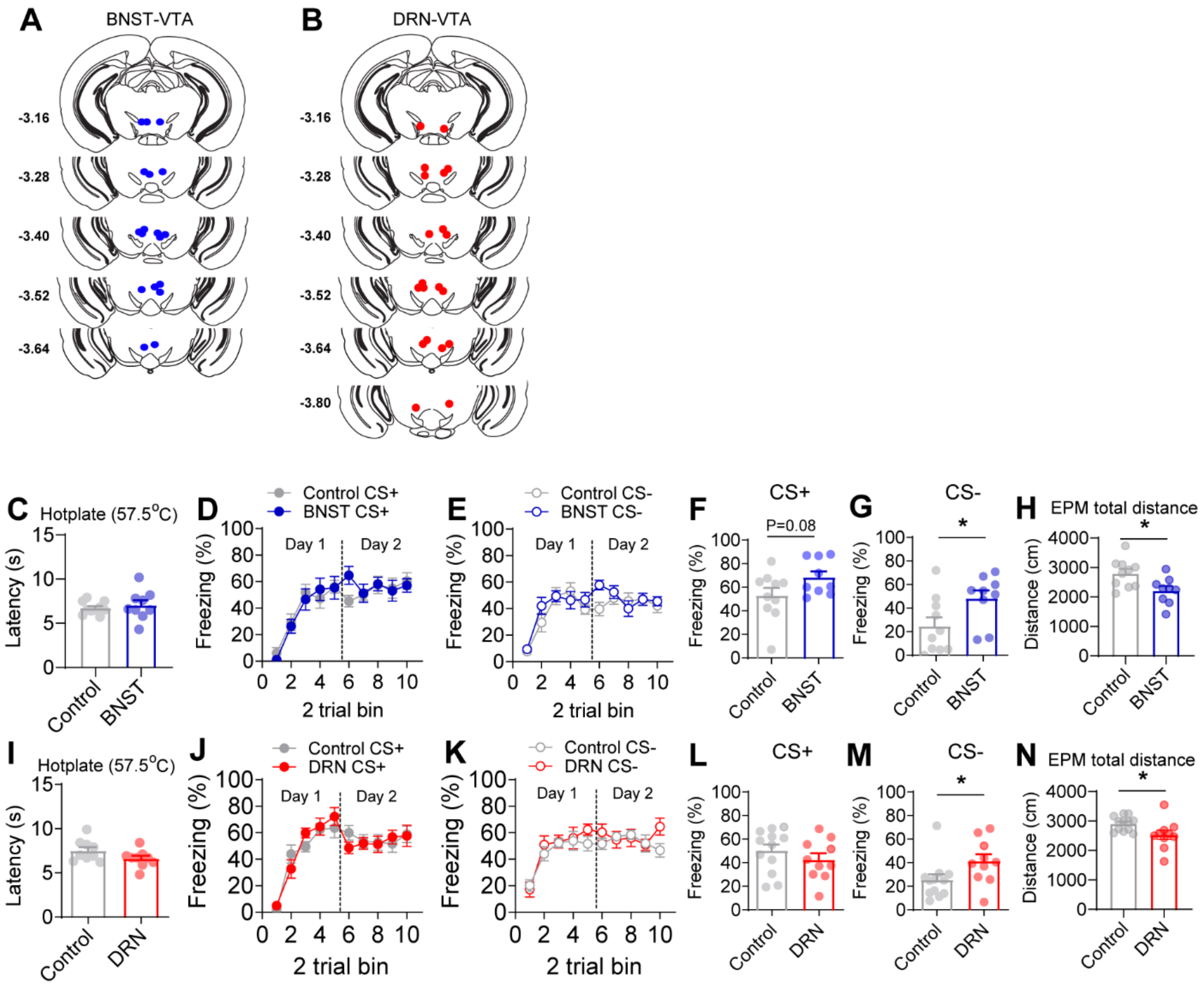


**Figure S4 related to Figures 3 and 4. Extended behavioral analysis of BNST→VTA and DRN→VTA ChR2-mCherry and control mice .** (A-B) Optical fiber placement in BNST→VTA (A) and DRN→VTA mice (B). (C) Latency to paw-lick or jump on a 57.5$^0$C hotplate in BNST→VTA or control *Pdyn*-Cre mice. (D-E) Average freezing behavior during conditioning days 1 and 2 in response to CS+ (D) and CS- (E) in BNST→VTA or control *Pdyn*-Cre mice. (F-G) Freezing to CS+ (F) or CS- (G) on test day in BNST→VTA or control *Pdyn*-Cre mice (F, Two-tailed Student's t-test, $t_{(17)}$=1.85, P=0.08; G, Two-tailed Student's t-test, $t_{(17)}$=2.29, P=0.035). (H) Total distance traveled during elevated plus maze assay in BNST→VTA or control *Pdyn*-Cre mice (Two-tailed Student's t-test, $t_{(17)}$=2.71, P=0.015). (C-H) N=10 control and N=9 BNST→VTA mice. (I) Latency to paw-lick or jump on a 57.5$^0$C hotplate in DRN→VTA or control *Pdyn*-Cre mice. (J-K) Average freezing behavior during conditioning days 1 and 2 in response to CS+ (J) and CS- (K) in DRN→VTA or control *Pdyn*-Cre mice. (L-M) Freezing to CS+ (L) or CS- (M) on test day in DRN→VTA or control *Pdyn*-Cre mice (M, Two-tailed Student's t-test, $t_{(20)}$=2.12, P=0.047). (N) Total distance traveled during elevated plus maze assay in DRN→VTA or control *Pdyn*-Cre mice (Two-tailed Student's t-test, $t_{(20)}$=2.34, P=0.03). (I-M) N=12 control and N=10 DRN→VTA mice. Data are presented as mean ± S.E.M.
